# The RNA m^6^A reader YTHDF1 promotes hematopoietic malignancy by maintaining oncogenic translation

**DOI:** 10.1101/2022.10.08.511406

**Authors:** Yun-Guang Hong, Zhigang Yang, Yan Chen, Tian Liu, Yuyuan Zheng, Guo-Cai Wu, Yinhui Chen, Juan Xia, Ruiting Wen, Wenxin Liu, Yi Zhao, Jin Chen, Xiangwei Gao, Zhanghui Chen

**Author notes:** These authors contributed equally (Yun-Guang Hong, Zhigang Yang). Correspondence: Zhanghui Chen, or Xiangwei Gao,.

## Abstract

N6-methyladenosine (m^6^A), the most abundant modification in mRNAs, has been defined as a crucial modulator in the progression of acute myeloid leukemia (AML), while the detailed mechanism remains elusive. Here we report that YTHDF1, an m^6^A reader protein, is overexpressed in human AML samples with enrichment in leukemia stem cells (LSCs). Whereas YTHDF1 is dispensable for normal hematopoiesis in mice, depletion of YTHDF1 attenuates self-renewal and proliferation of patient-derived LSCs, and impedes leukemia establishment in immunodeficient mice. Mechanistically, YTHDF1 promotes the translation of diverse m^6^A-modified oncogene mRNAs particularly *cyclin E2*. We applied a structure-based virtual screening of FDA-approved drugs and identified tegaserod as a potential YTHDF1 inhibitor. Tegaserod blocks the direct binding of YTHDF1 with m^6^A-modified mRNAs and inhibits YTHDF1-regulated mRNA translation. Moreover, tegaserod inhibits leukemogenesis *in vitro* and in mice, phenocopying the loss of YTHDF1. Together, our study defines YTHDF1 as an integral regulator of AML progression at the translational level and identifies tegaserod as a potential therapeutic agent for AML by targeting YTHDF1.

**Statement of Significance:**

1. The RNA m^6^A reader YTHDF1 is an oncoprotein in AML.
2. YTHDF1 promotes the translation of m^6^A-modified oncogene mRNAs.
3. Tegaserod, an FDA-approved drug for irritable bowel syndrome, inhibits YTHDF1-regulated mRNA translation and leukemogenesis.

**Graphical Abstract:** 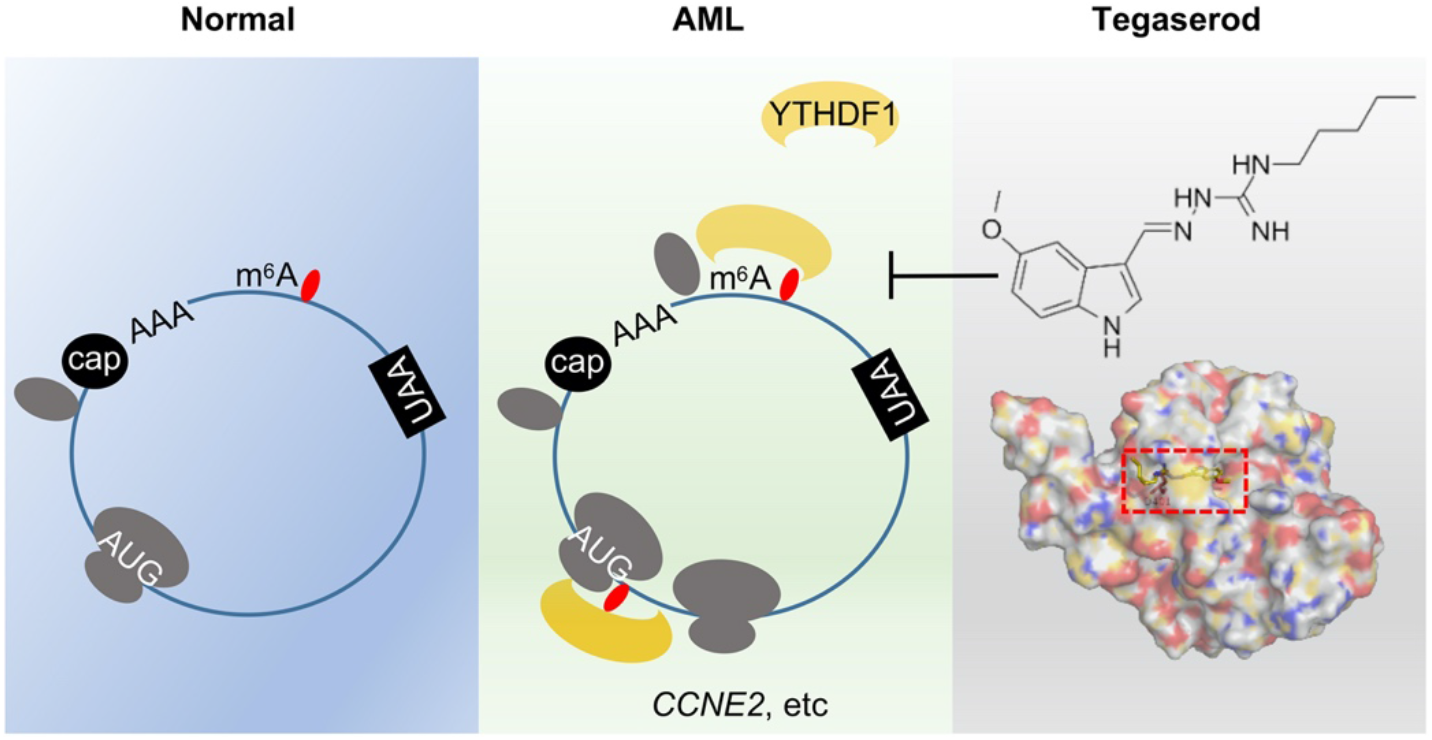

## INTRODUCTION

Acute myeloid leukemia (AML) is a heterogeneous hematologic malignancy characterized by impaired differentiation and uncontrolled proliferation of immature myeloid cells (1,2). As the most common form of acute leukemia among adults, AML accounts for the largest number of annual deaths from leukemias in the world. With available therapeutics, only less than 30% of AML patients can survive more than 5 years (3). More effective targeted therapies are urgently needed for human AML treatment, which relies on a better understanding of molecular mechanisms underlying the pathogenesis of AML.

Although cytogenetic abnormalities, including balanced translocations and chromosomal gains or losses, play an integral role in the development of AML, accumulating evidence has highlighted the essential role of dysregulated epigenetic processes in the pathogenesis of AML (4). Genes involved in DNA modifications or histone modifications, such as *DNMT3A, KMT2A*, and *TET2*, show recurrent somatic alterations or altered expression in AML (5). These epigenetic alterations induce dysregulation of gene expression without modifying the DNA coding sequence, leading to the development of AML. Specific inhibitors targeting epigenetic modifications have shown promising therapeutic potential for AML (5). However, epigenetic dysregulation in AML is more common than expected, and additional layers of epigenetic alterations remain to be elucidated.

N6-methyladenosine, the most abundant modification on mRNAs, is emerging as a new layer of gene expression regulation. The formation of m^6^A is catalyzed by RNA methyltransferase complex METTL3/METTL14/WTAP (6–8), while demethylation is mediated by ALKBH5 or FTO (9,10). The YTH domain family proteins serve as m^6^A “reader” proteins to regulate a diverse array of RNA metabolism, including mRNA splicing, degradation, and translation (11,12). m^6^A modification has been defined as a crucial modulator in AML progression, while the overall impact of this modification is complex. The demethylases FTO and ALKBH5 have been reported to promote the tumorigenicity of AML (13,14). However, METTL3 and METTL14, two components of the methyltransferase complex, are also critical promoters for AML pathogenesis (15–17). Since the fate of the methylated mRNAs, ranging from degradation to translation, is decided by its reader proteins (11,12), we proposed that deciphering the function of m^6^A reader proteins will help elucidate the complexity of m^6^A in AML.

In this study, we demonstrated an essential role of YTHDF1 in AML development and maintenance. Genome-wide analysis revealed a subset of oncogenes regulated by YTHDF1 including *CCNE2*. Importantly, we identified tegaserod, an FDA-approved drug for irritable bowel syndrome with constipation (18), as a potential therapeutic agent for AML through blocking YTHDF1’s function. To our knowledge, this is the first candidate drug targeting YTHDF1-regulated mRNA translation in cancer.

## RESULTS

### The protein level of YTHDF1 is highly expressed in AML

To explore the protein level of YTHDF1 in AML, we detected the expression of YTHDF1 protein in human AML patients by using flow cytometry. Data revealed that the expression of YTHDF1 was significantly up-regulated in human AML samples than the non-leukemic control group (Fig. 1a-b). We further classified AML samples according to the French-American-British (FAB) classification system of acute leukemia and analyzed YTHDF1 expression (19). All of the AML samples with distinct FAB subtypes showed higher YTHDF1 expression than control group, while no significant difference was observed among different FAB subtypes (Fig. 1c). Consistent with the flow cytometry data, the immunoblotting analysis showed increased YTHDF1 protein level in AML samples (Fig. 1d). These results indicated that YTHDF1 protein is elevated in AML samples. LSCs are considered to be the root cause for AML progression (20). We therefore isolated CD34^+^ LSCs from the primary AML patient samples and determined YTHDF1 protein level. Strikingly, the YTHDF1 protein level in CD34^+^ LSCs is ubiquitously higher than the level in normal cord blood-derived CD34^+^ cells and CD34^-^ AML cells (Fig. 1e-h), implying a role of this protein in LSCs function.

**Figure 1.**
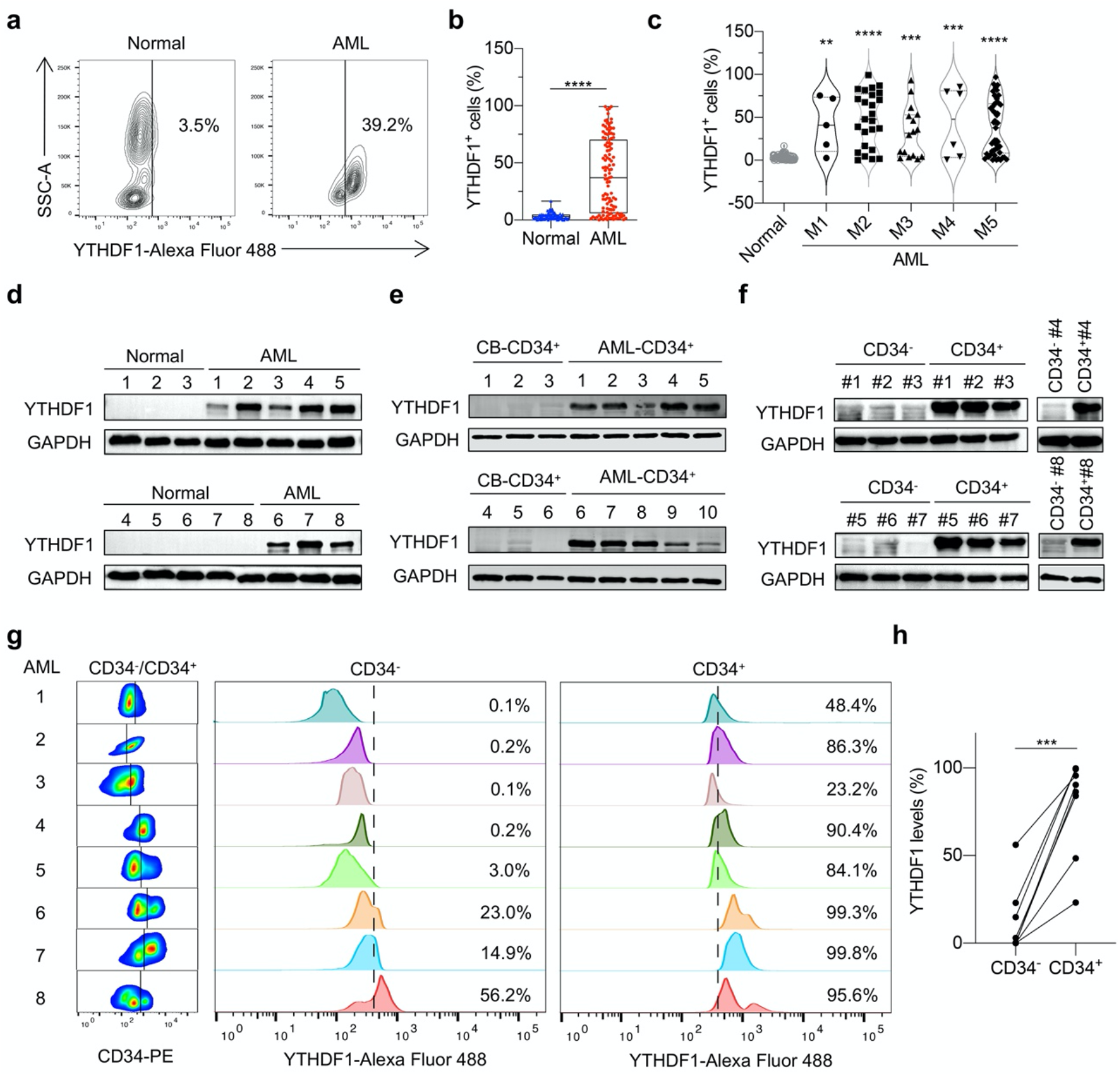
The protein level of YTHDF1 is highly expressed in AML. (a) Representative contour map of YTHDF1 staining from one healthy donor (Normal) blood sample and one AML sample. (b) Percentage of YTHDF1^+^ cells in Normal (n = 80) and AML (n = 115) samples. Data are represented as mean ± SEM. \*\*\*\**p* < 0.0001, t-test. (c) Violin plot showing the percentage of YTHDF1^+^ cells in distinct FAB subtypes of AML samples. ** *p* < 0.01, *** *p* < 0.001, \*\*\*\**p* < 0.0001, one-way ANOVA. (d) Immunoblot analysis of YTHDF1 expression in peripheral blood mononuclear cells (PBMCs) from Normal (n = 8) and AML samples (n = 8). GAPDH was used as a loading control. (e) Immunoblot analysis of YTHDF1 expression in normal cord blood-derived CD34^+^ cells (n = 6) and AML CD34^+^ LSCs (n = 10). GAPDH was used as a loading control. (f) Immunoblot analysis of YTHDF1 expression in patient-derived CD34^+^ LSCs and CD34^-^ AML cells. GAPDH was used as a loading control. (g) YTHDF1 abundance in patient-derived CD34^+^ LSCs and CD34^-^ AML cells. (h) Statistical analysis of YTHDF1^+^ cells in patient-derived CD34^+^ LSCs versus CD34^-^ AML cells in (g) (n = 8). *** *p* < 0.001, t-test.

However, the Cancer Genome Atlas (TCGA) data (Supplementary Fig. 1a), GSE data (GSE83533 and GSE183817, Supplementary Fig. 1b), and our qPCR data (Supplementary Fig. 1c-d) showed no difference between normal human and AML patients. We concluded that the expression of YTHDF1 is regulated at protein level, but not mRNA level in AML.

### Depletion of *Ythdf1* gene does not alter hematopoiesis in mice

We explored the impact of *Ythdf1* knockout on hematopoietic stem and progenitor cells in mice. Data showed that the adult *Ythdf1^-/-^* mice displayed unaffected proportions of Lin^-^, Sca-1^+^ or/and c-Kit^+^ cells in bone marrow (BM) compared to wild-type (WT) mice (Fig. 2a), suggesting that YTHDF1 does not affect steady-state hematopoiesis. We further investigated whether *Ythdf1* deletion affects the differentiation of normal hematopoietic stem cells and multilineage hematopoiesis. To this end, differentiated cells in peripheral blood (PB) were determined by using myeloid cell markers (Gr-1 and CD11b) and lymphoid cell markers (CD3, CD4, and CD19). There was no significant difference in the proportion of terminal differentiated myeloid cells, B lymphocytes, and T lymphocytes between *Ythdf1^-/-^* mice and WT mice (Fig. 2b-c). Compared with WT mice, *Ythdf1^-/-^* mice showed similar blood cell counts and proportion of different types of blood cells in the PB, including white blood cells (WBCs), lymphocytes (LYMs), neutrophils (NEUs), monocytes (MONs), platelets (PLT), red blood cells (RBCs), and hemoglobin (HGB) (Fig. 2d).

**Figure 2.**
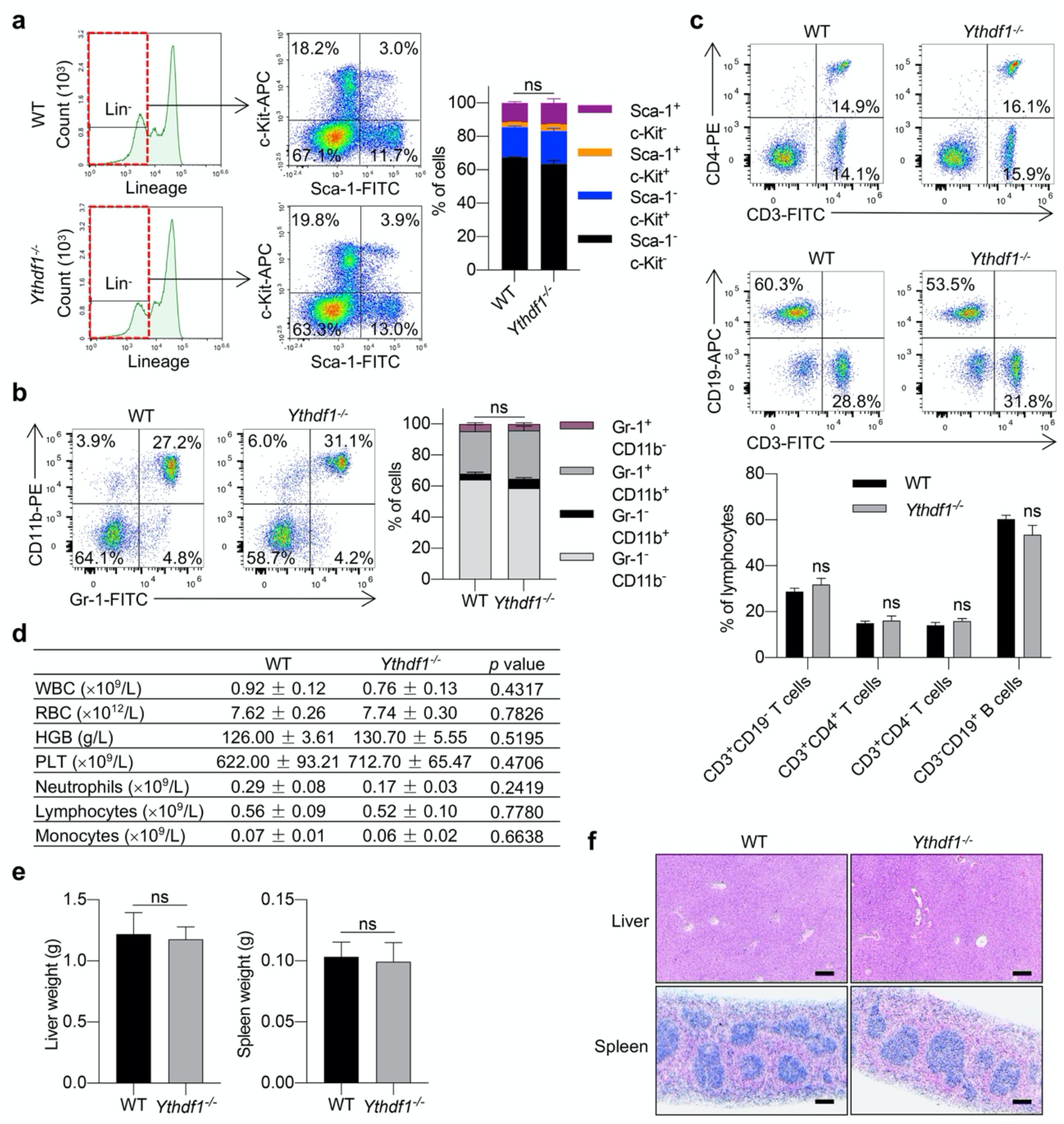
Depletion of *Ythdf1* gene does not alter hematopoiesis in mice. (a) Representative flow cytometry plot of bone morrow (BM) Sca-1^+^ or/and c-Kit^+^ cell populations from *Ythdf1^-/-^* mice and WT mice. The quantitative data (right panel) are represented as mean ± SEM (n = 5, ns, not significant, two-way ANOVA). (b) Representative flow cytometry plot of peripheral blood (PB) terminal differentiated myeloid cell populations from *Ylhdf1^-/-^* mice and WT mice. The quantitative data (right panel) are represented as mean ± SEM (n = 5, ns, not significant, two-way ANOVA). (c) Representative flow cytometry plot of terminal differentiated B and T lymphocyte subpopulations from *Ythdf1^-/-^* mice and WT mice. The quantitative data (bottom) are represented as mean ± SEM (n = 5, ns, not significant, two-way ANOVA). (d) Peripheral blood counts in *Ythdf1^-/-^* mice and WT mice. Data are represented as mean ± SEM (n = 4, t-test). (e) The weight of liver and spleen from *Ythdf1^-/-^* mice and WT mice (n = 3, ns, not significant, t-test). (f) Hematoxylin and eosin (H&E) staining of liver and spleen from *Ythdf1^-/-^* mice and WT mice. Scale bar, 100 μm.

We also detected the development of extramedullary hematopoietic organs in *Ythdf1^-/-^* mice. The weight of liver and spleen was similar between *Ythdf1^-/-^* mice and WT mice (Fig. 2e). Moreover, the organ architecture of liver and spleen remained normal in these mice (Fig. 2f). Taken together, depletion of *Ythdf1* has little effect on the differentiation of normal hematopoietic stem cells and the development of extramedullary hematopoietic organs.

### YTHDF1 is essential for the progression of AML

To study the function of YTHDF1 in human AML development, we silenced YTHDF1 using two independent shRNAs targeting YTHDF1 (21) (hereafter referred to as shDF1#1, shDF1#2), which showed high efficiency in THP-1 and MV-4-11 human AML cells (Fig. 3a). YTHDF1 knockdown (KD) significantly inhibited cell proliferation, induced G1 phase arrest, and promoted cell apoptosis of THP-1 and MV-4-11 cells (Fig. 3b-d). We further measured the colony-forming capacity of THP-1 and MV-4-11 cells. YTHDF1 KD group had significantly fewer colonies formed than the control group (Fig. 3e-f, Supplementary Fig. 2a), indicating that LSCs function was significantly impaired following YTHDF1 KD.

**Figure 3.**
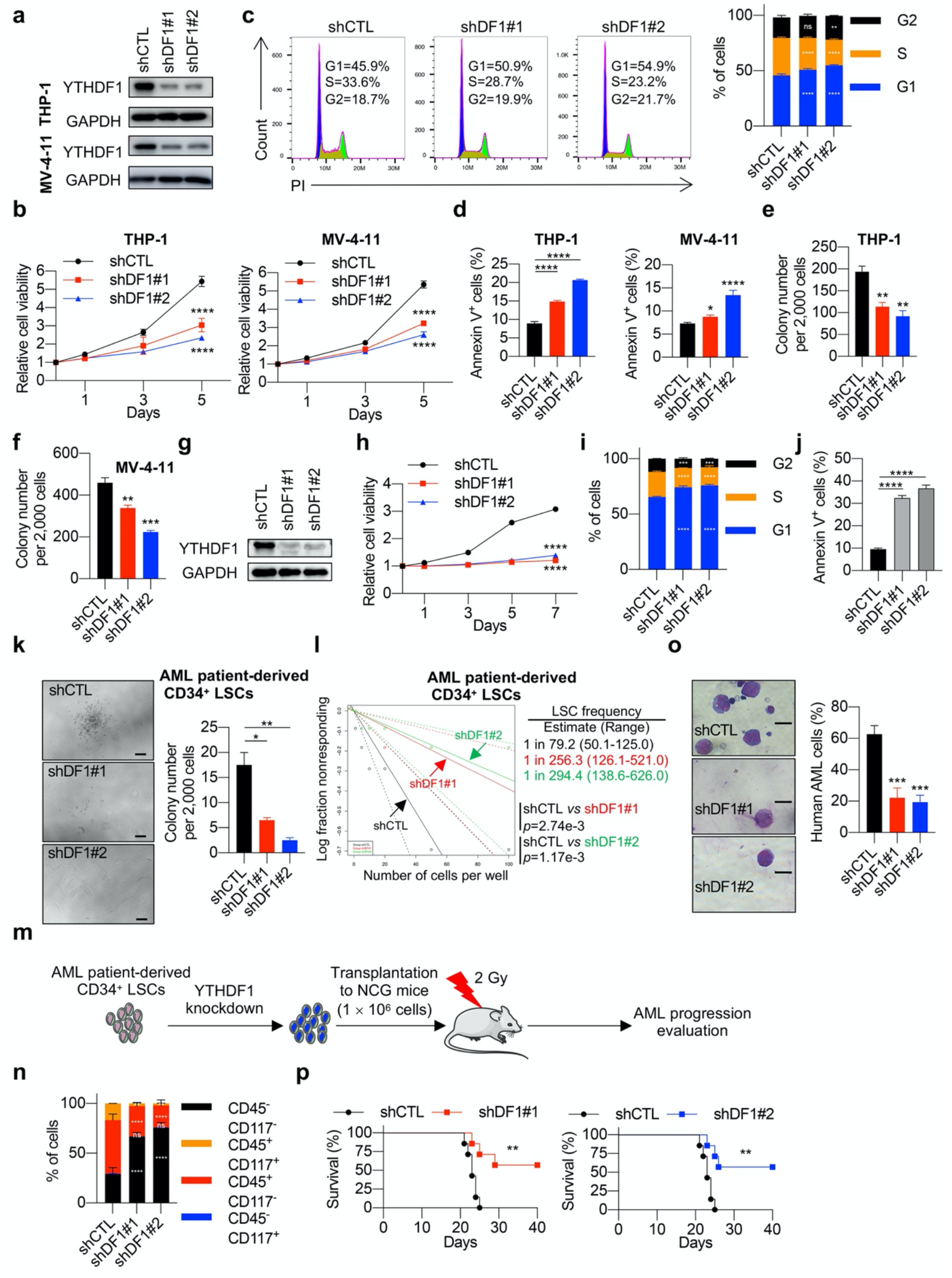
YTHDF1 is essential for the progression of AML. (a) Immunoblot analysis of YTHDF1 in control and YTHDF1 knockdown THP-1 and MV-4-11 cells. GAPDH was used as a loading control. (b) Cell proliferation analysis in control and YTHDF1 knockdown THP-1 and MV-4-11 cells. Data are represented as mean ± SEM (n = 3). **** *p* < 0.0001, two-way ANOVA. (c) Cell cycle analysis in control and YTHDF1 knockdown THP-1 cells. The quantitative data (right panel) are represented as mean ± SEM (n = 3). ** *p* < 0.01, \*\*\*\**p* < 0.0001, ns, not significant, two-way ANOVA. (d) Annexin V staining of control and YTHDF1 knockdown THP-1 and MV-4-11 cells. The quantification of Annexin V^+^ cells are represented as mean ± SEM (n = 3). * *p* < 0.05, **** *p* < 0.0001, t-test. (e-f) Colony-forming units of control and YTHDF1 knockdown THP-1 and MV-4-11 cells. The quantitative data are represented as mean ± SEM (n = 3). \*\**p* < 0.01, *** *p* < 0.001, t-test. (g) Immunoblot analysis of YTHDF1 in AML patient-derived CD34^+^ LSCs with or without YTHDF1 knockdown. GAPDH was used as a loading control. (h) Cell proliferation analysis in AML patient-derived CD34^+^ LSCs with or without YTHDF1 knockdown. Data are represented as mean ± SEM (n = 3). \*\*\*\**p* < 0.0001, two-way ANOVA. (i) Cell cycle analysis in AML patient-derived CD34^+^ LSCs with or without YTHDF1 knockdown. The quantitative data are represented as mean ± SEM (n = 3). \*\*\**p* < 0.001, **** *p* < 0.0001, two-way ANOVA. (j) Annexin V staining of AML patient-derived CD34^+^ LSCs with or without YTHDF1 knockdown. The quantification of Annexin V^+^ cells are represented as mean ± SEM (n = 3). **** *p* < 0.0001, t-test. (k) Colony-forming units of AML patient-derived CD34^+^ LSCs with or without YTHDF1 knockdown. The quantitative data (right panel) are represented as mean ± SEM (n = 3). * *p* < 0.05, ** *p* < 0.01, t-test. Scale bar, 200 μm. (l) LSC frequency changes in AML patient-derived CD34^+^ LSCs upon YTHDF1 KD as estimated by *in vitro* limiting dilution assays (LDAs). (m) Schematic of xenograft mouse model. Irradiated immunodeficient mice (NCG) were injected with control or YTHDF1 knockdown AML patient-derived CD34^+^ LSCs and analyzed for AML progression. (n) Percentage of human CD45^+^ and CD117^+^ cells in the bone morrow of recipient mice. The quantitative data are represented as mean ± SEM (n = 3). ** *p* < 0.01, **** *p* < 0.0001, ns, not significant, two-way ANOVA. (o) Wright-Giemsa staining of bone morrow from recipient mice. The quantitative data (right panel) are represented as mean ± SEM (n = 3). \*\*\**p* < 0.001, t-test. Scale bar, 10 μm. (p) Survival curve of NCG mice transplanted with control or YTHDF1 knockdown AML patient-derived CD34^+^ LSCs (7 mice per group). ** *p* < 0.01, Mantel-Cox test.

To consolidate the results in THP-1 and MV-4-11 cells, we performed these experiments in AML patient-derived CD34^+^ LSCs. YTHDF1 KD substantially inhibited cell proliferation, induced G1 phase arrest, and promoted cell apoptosis (Fig. 3g-j). Moreover, YTHDF1 KD suppressed the colony-forming capability of CD34^+^ LSCs (Fig. 3k). Limiting dilution assays further demonstrated that YTHDF1 KD resulted in a remarkable decrease in the frequency of CD34^+^ LSCs (Fig. 3l).

To evaluate the role of YTHDF1 in leukemogenesis *in vivo*, AML patient-derived CD34^+^ LSCs expressing a scrambled shRNA or YTHDF1 shRNAs were transplanted into immunodeficient recipient mice via tail vein injection (Fig. 3m). Compared with control cells, YTHDF1-deficient cells had lower engrafting capacity into the bone morrow (Fig. 3n-o, Supplementary Fig. 2b). The body weight was reduced in mice transplanted with control AML cells but did not change in the YTHDF1 knockdown group (Supplementary Fig. 2c). Knockdown of YTHDF1 also significantly reduced the AML cells infiltration of liver architecture (Supplementary Fig. 2d). Importantly, YTHDF1 knockdown markedly prolonged the survival of recipient mice (Fig. 3p). Together, these data indicated that YTHDF1 promotes AML progression.

### Transcriptome-wide identification of YTHDF1-regulated transcripts

YTHDF1 is a well-known m^6^A reader protein that regulates gene expression at the translational level. To identify YTHDF1-regulated transcripts, we performed m^6^A-seq in THP-1 cells. In line with previous studies, m^6^A-seq revealed the expected distribution of m^6^A peaks enriched at the stop codon within transcripts (Fig. 4a). The m^6^A peaks were characterized by the canonical RGACH motif (Fig. 4b). To analyze mRNA and translational changes, we performed RNA sequencing (RNA-seq) and ribosome profiling (Ribo-seq) in THP-1 cells after YTHDF1 knockdown. We detected 1,006 decreased genes and 1,293 increased genes from Ribo-seq data, but slight changes from RNA-seq data in YTHDF1-deficient cells (Fig. 4c and Supplementary Table 2). By overlapping with m^6^A-modified mRNAs, we identified 531 decreased genes and 664 increased genes at the translational level (Fig. 4d). Kyoto Encyclopedia of Genes and Genomes (KEGG) pathway analysis of the 531 decreased genes showed that a group of oncogenic pathways was enriched. The top significantly enriched pathway was the PI3K-AKT signaling pathway (Fig. 4e and Supplementary Table 3), which contributes to AML progression by regulating cell proliferation, apoptosis, and differentiation (22). In addition to the downregulated genes, we further analyzed the 664 upregulated genes using KEGG pathway analysis. The results showed the top significantly enriched pathway was Endocytosis (Supplementary Fig. 3a). Together, these data supported an oncogenic role of YTHDF1 in AML development.

**Figure 4.**
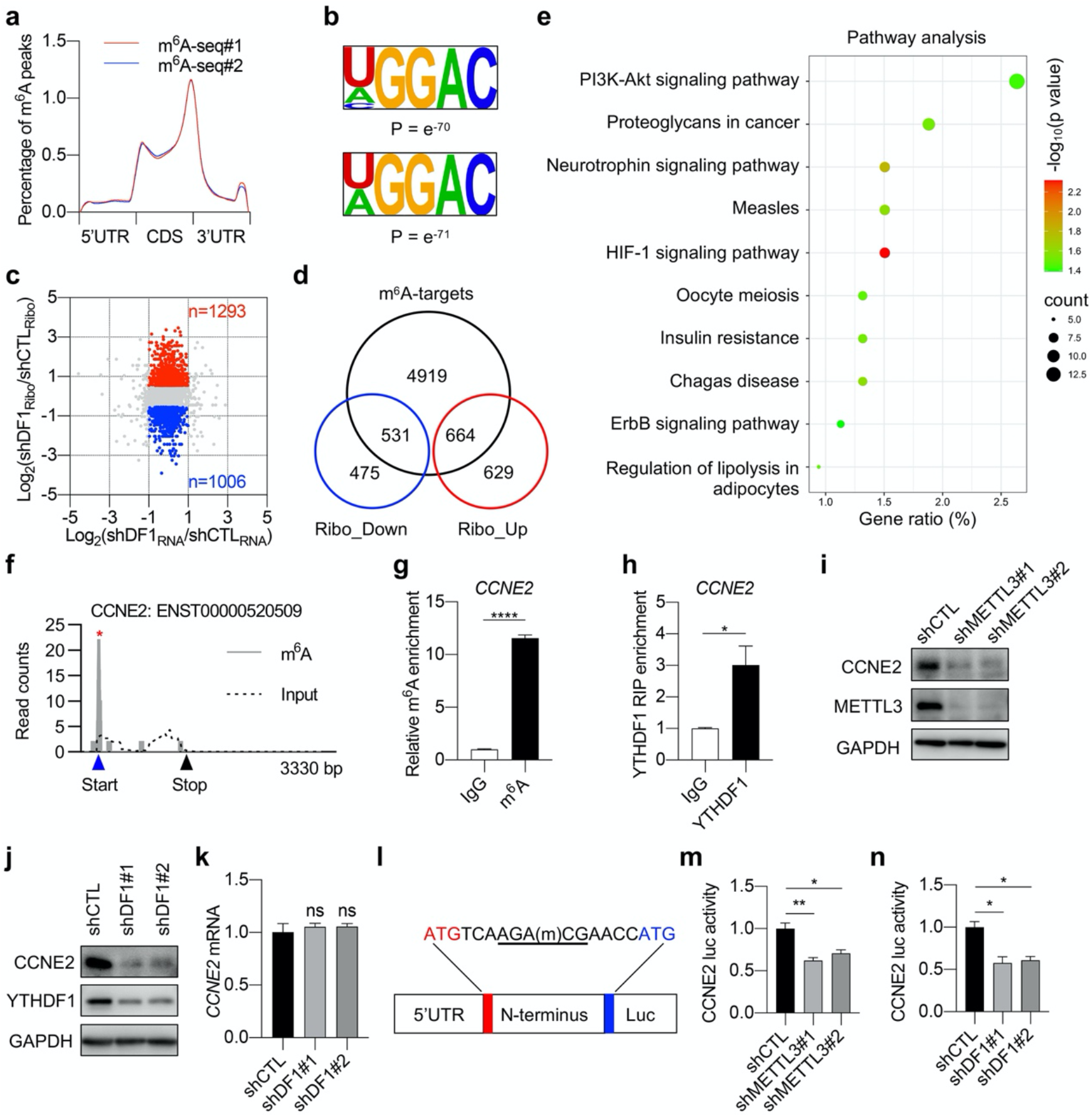
YTHDF1 regulates the translation of *CCNE2* in AML. (a) Metagene profiles of m^6^A distribution across transcripts in THP-1 cells. (b) Consensus motif of m^6^A sites in THP-1 cells. (c) Scatterplot of RNA-seq data and ribosome profiling data from control and YTHDF1 knockdown cells. The upregulated (red) and downregulated (blue) genes at the translational level are highlighted. (d) The overlap of m^6^A-modified mRNAs with differential expressed mRNAs at the translational level. (e) KEGG enrichment analysis of YTHDF1 targets that are downregulated in YTHDF1-deficient cells. (f) The m^6^A distribution in *CCNE2* transcript. * indicates the predicated m^6^A peak. (g) Me-RIP-qPCR analysis of the m^6^A level of *CCNE2*. Data are represented as mean ± SEM (n = 3). **** *p* < 0.0001, t-test. (h) RIP-qPCR analysis of the binding of YTHDF1 with *CCNE2* mRNA. Data are represented as mean ± SEM (n = 3). * *p* < 0.05, t-test. (i) Immunoblot analysis of CCNE2 in control and METTL3 knockdown THP-1 cells. (j) Immunoblot analysis of CCNE2 in control and YTHDF1 knockdown THP-1 cells. (k) Relative *CCNE2* mRNA level in control and YTHDF1 knockdown THP-1 cells (n = 3, ns, not significant, t-test). (l) Schematic of the construct used for luciferase assay. (m) Luciferase activity of the construct in (l) in control and METTL3 knockdown cells. Data are represented as mean ± SEM. * *p* < 0. 05, ** *p* < 0. 01 (n = 3, t-test). (n) Luciferase activity of the construct in (l) in control and YTHDF1 knockdown cells. Data are represented as mean ± SEM. * *p* < 0. 05 (n = 3, t-test).

### YTHDF1 regulates *CCNE2* translation in an m^6^A-dependent manner

Focusing on potential candidates implicated in PI3K-AKT signaling, cyclin E2 (CCNE2) was most significantly downregulated listed as the candidate upon YTHDF1 deletion (Supplementary Fig. 3b), we noticed that *cyclin E2* (*CCNE2*), bearing an m^6^A site downstream the start codon, showed reduced translational efficiency upon YTHDF1 deletion (Fig. 4f). Methylated RNA immunoprecipitation (Me-RIP) confirmed the presence of m^6^A in *CCNE2* mRNA (Fig. 4g). RNA immunoprecipitation (RIP) confirmed the direct binding between YTHDF1 and *CCNE2* mRNA (Fig. 4h). Knockdown of METTL3 reduced the protein level of CCNE2 (Fig. 4i), implying a role of m^6^A in regulating CCNE2 expression. YTHDF1 knockdown dramatically reduced the protein level but not the mRNA level of CCNE2 (Fig. 4j-k), suggesting that the regulation happens at the translational level. Finally, we investigated the regulation of *CCNE2* translation by m^6^A/YTHDF1 using a luciferase assay. The 5’UTR together with the CDS region containing the predicted m^6^A site was fused in frame with luciferase gene (Fig. 4l). Erasing m^6^A by silencing METTL3 decreased *CCNE2* luciferase activity (Fig. 4m), suggesting the involvement of m^6^A in *CCNE2* translation. YTHDF1 knockdown decreased luciferase activity (Fig. 4n), suggesting that YTHDF1 mediates the translation of *CCNE2*. Collectively, these results indicated that YTHDF1 promotes the translation of *CCNE2* in an m^6^A-dependent manner.

### CCNE2 plays an oncogenic role in AML

Both the mRNA level and the protein level of CCNE2 were elevated in human AML samples (Fig. 5a-b). TCGA database also revealed elevated expression of *CCNE2* in AML samples, implying an oncogenic role (Fig. 5c). Knockdown of CCNE2 dramatically reduced cell proliferation and S phase transition, and induced cell apoptosis (Fig. 5d-g). Next, we performed rescue experiments by overexpressing CCNE2 in YTHDF1-deficient THP-1 cells (Fig. 5h). Cell proliferation and cell cycle suppressed by YTHDF1 deficiency were re-established after CCNE2 overexpression (Fig. 5i-j). In addition, overexpression of CCNE2 reversed the increased cell apoptosis by YTHDF1 knockdown (Fig. 5k). These results suggested that CCNE2 is a functional target of YTHDF1 to facilitate AML progression.

**Figure 5.**
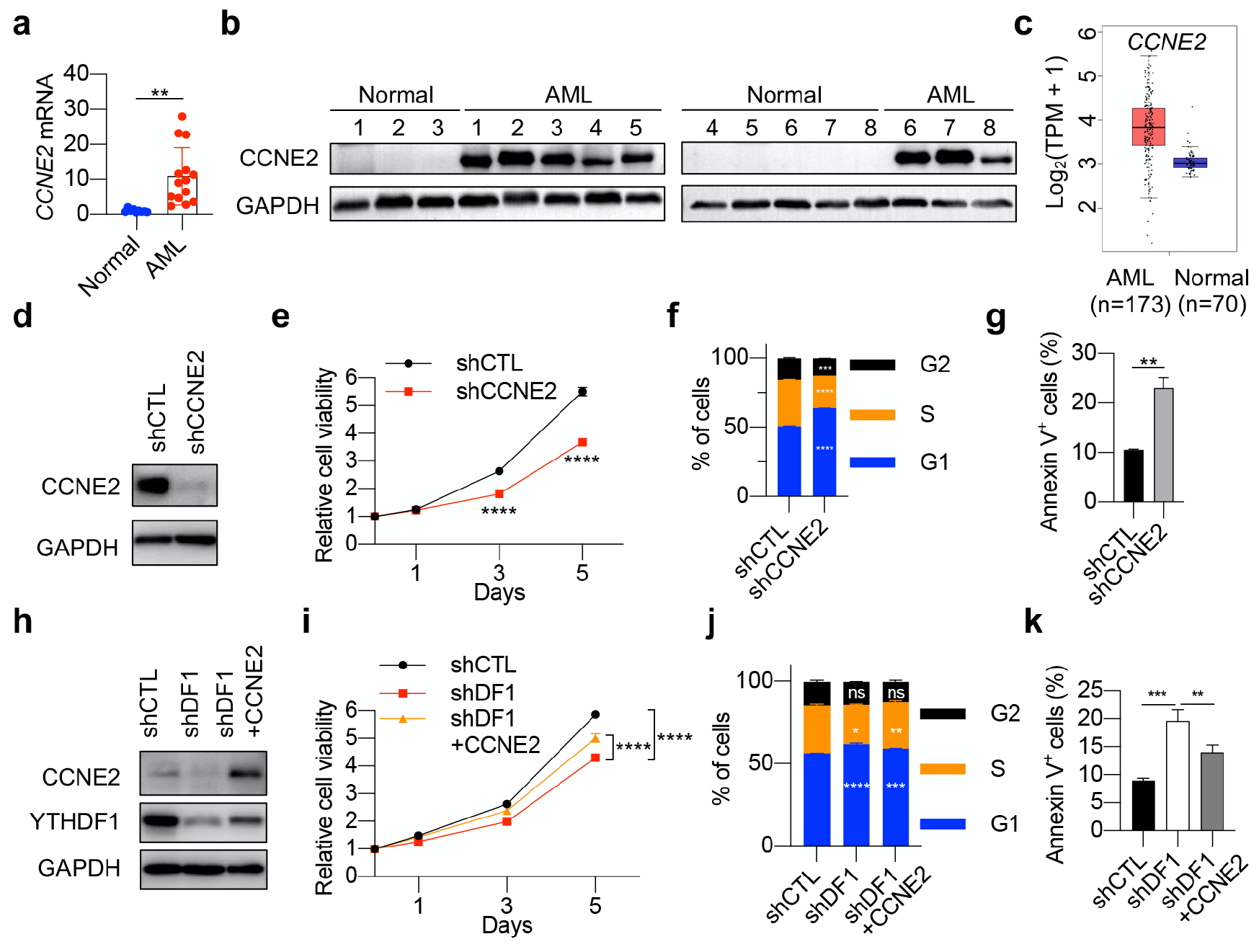
CCNE2 plays an oncogenic role in AML. (a) Relative *CCNE2* mRNA level in Normal (n = 9) and AML samples (n = 14). Data are represented as mean ± SEM. \*\**p* < 0.01, t-test. (b) Immunoblot analysis of CCNE2 in PBMCs from Normal (n = 8) and AML samples (n = 8). GAPDH was used as a loading control. (c) Box plot showing the relative *CCNE2* mRNA expression in Normal and AML samples from the TCGA database. (d) Immunoblot analysis of CCNE2 in control and CCNE2 knockdown THP-1 cells. GAPDH was used as a loading control. (e) Cell proliferation of control and CCNE2 knockdown THP-1 cells. Data are represented as mean ± SEM (n = 3). \*\*\*\**p* < 0.0001, two-way ANOVA. (f) Cell cycle analysis in control and CCNE2 knockdown THP-1 cells. The quantitative data are represented as mean ± SEM (n = 3). *** *p* < 0.001, \*\*\*\**p* < 0.0001, two-way ANOVA. (g) Annexin V staining of control and CCNE2 knockdown THP-1 cells. The quantification of Annexin V^+^ cells are represented as mean ± SEM (n = 3). ** *p* < 0.01, t-test. (h) Immunoblot analysis of YTHDF1 and CCNE2 in YTHDF1-deficient THP-1 cells upon overexpression of CCNE2. (i) Cell proliferation was measured in THP-1 cells described in (h). Data are represented as mean ± SEM. **** *p* < 0.0001, two-way ANOVA. (j-k) Cell cycle (j) and apoptosis (k) were determined by flow cytometry in THP-1 cells described in (h). Data are represented as mean ± SEM (n = 3). shDF1 group was compared with shCTL group. shDF1+CCNE2 group was compared with shDF1 group. * *p* < 0.05, ** *p* < 0.01, *** *p* < 0.001, \*\*\*\**p* < 0.0001, ns, not significant. Two-way ANOVA was used for cell cycle (j) analysis, while t-test (k) was used for apoptosis analysis.

### Tegaserod blocks the binding of YTHDF1 with m^6^A-modified mRNAs

YTHDF1 is dispensable for AML development but not required for normal hematopoiesis, making it an ideal target for AML treatment. To discover specific pharmacological inhibitors targeting YTHDF1-m^6^A binding, we performed molecular docking based on the crystal structure of YTH domain of YTHDF1 (Fig. 6c) (23). From an FDA-approved drug library (totally 1805 compounds), we identified 21 compounds with high docking scores (Fig. 6a and Supplementary Table 4). Among them, tegaserod showed high binding energy with the YTH domain (−4.725 kcal/mol) and the most dramatic inhibition on THP-1 cell proliferation (Fig. 6b). In the binding model, tegaserod forms three hydrogen bonds with the residues TRP470, TYR397, and CYS412 and two π-π interactions with TYR397 and TRP470 (Fig. 6c), overlapping with the m^6^A-binding region. Moreover, we further verified that tegaserod could directly bind to the YTH domain of YTHDF1 in a concentration-dependent manner via a surface plasmon resonance (SPR) binding assay (Supplementary Fig. 4a).

**Figure 6.**
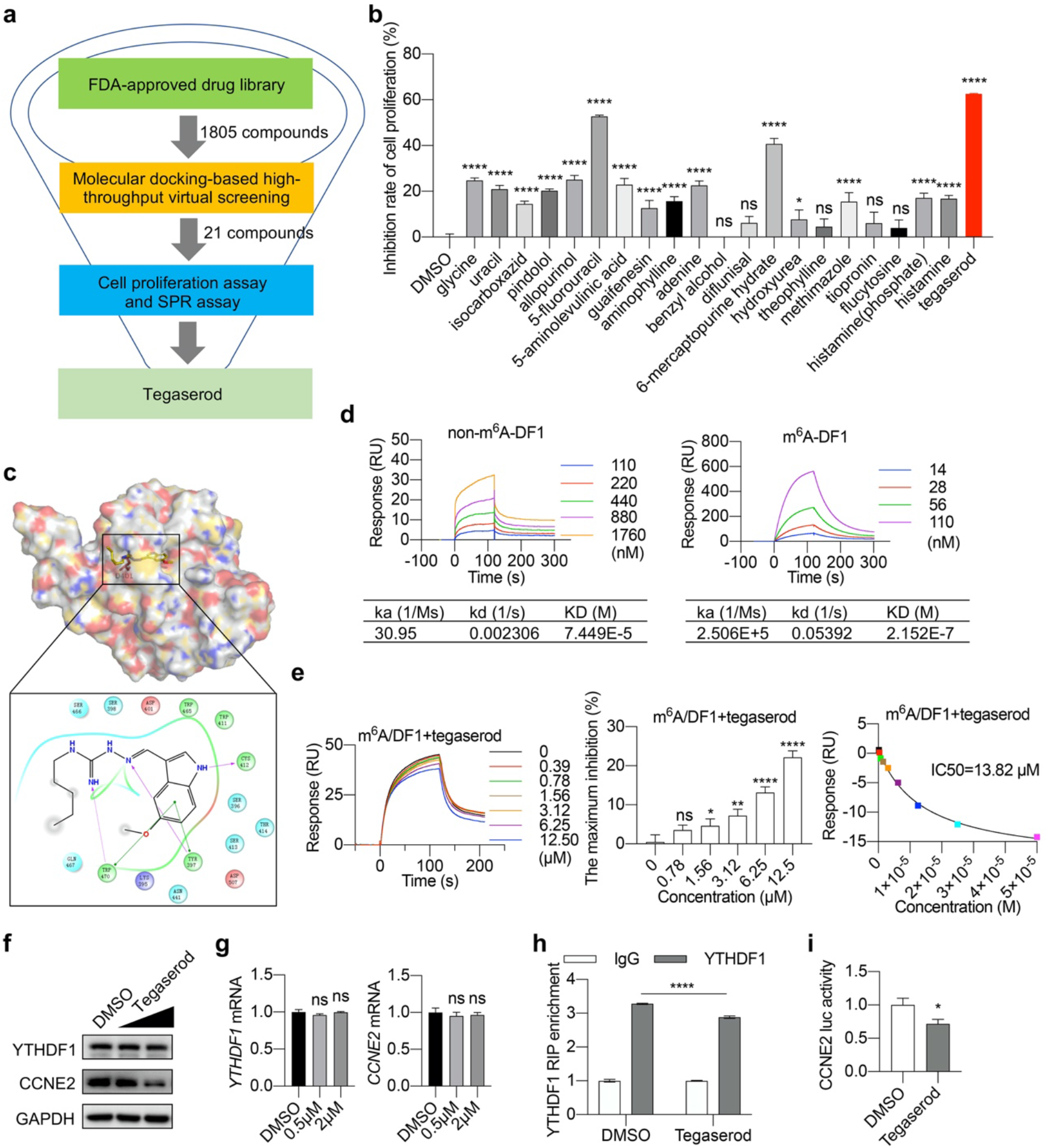
Tegaserod blocks the binding of YTHDF1 to m^6^A-modified mRNAs. (a) Flowchart of structure-based virtual screening. (b) Effect of 21 pharmacological chemicals on the viability of THP-1 cells (n = 3, \**p* < 0.05, \*\*\*\**p* < 0.0001, ns, not significant, t-test). (c) Molecular docking of tegaserod interacting with YTH domain of YTHDF1. Tegaserod forms three hydrogen bonds with residues TRP470, TYR397, and CYS412 and two π-π interactions with TYR397 and TRP470. (d) Biacore assay of the binding between YTH domain of YTHDF1 and m^6^A-modified oligos or non-m^6^A-modified oligos. (e) Effect of tegaserod on the interaction between m^6^A-modified oligos and YTH domain of YTHDF1. * *p* < 0.05, \*\**p* < 0.01, \*\*\*\**p* < 0.0001, ns, not significant, t-test. (f) Immunoblot analysis of YTHDF1 and CCNE2 in THP-1 cells treated with or without tegaserod maleate. (g) Relative mRNA level of *CCNE2* and *YTHDF1* in THP-1 cells treated with or without tegaserod maleate. ns, not significant, t-test. (h) RIP-qPCR analysis of the binding of YTHDF1 with *CCNE2* mRNA in THP-1 cells treated with or without tegaserod maleate. Data are represented as mean ± SEM (n = 3). **** *p* < 0.0001, t-test. (i) Luciferase activity of CCNE2 reporter in cells treated with or without tegaserod maleate. Data are represented as mean ± SEM. \**p* < 0. 05 (n = 3, t-test).

To study whether tegaserod blocks the binding of YTHDF1 protein with m^6^A-modified mRNAs directly, SPR binding assay was conducted. Firstly, we compared the binding of the YTH domain of YTHDF1 with m^6^A-modified mRNAs and non-m^6^A-modified mRNAs. The *K_D_* values of the two groups were 0.2152 μM and 74.49 μM, respectively (Fig. 6d), supporting that YTHDF1 is an m^6^A-binding protein. Next, we immobilized m^6^A-modified mRNAs on the chip surface, and series concentration of tegaserod was mixed with the YTH domain of YTHDF1 (20 nM) to determine its effect on YTH domain/m^6^A binding. Data showed that tegaserod blocks m^6^A/YTHDF1 binding in a concentration-dependent manner, and the half-maximal inhibitory concentration (IC50) was 13.82 μM (Fig. 6e). However, tegaserod has little effect on the binding of m^6^A/YTHDF2 or m^6^A/YTHDF3 (Supplementary Fig. 4b-c). These data suggested the specificity of tegaserod on m^6^A/YTHDF1 binding.

Finally, we asked whether tegaserod affects the translation of YTHDF1 target mRNAs in THP-1 cells. Indeed, tegaserod treatment downregulated CCNE2 protein level without affecting its mRNA level, phenocopied YTHDF1 knockdown (Fig. 6f-g). RNA immunoprecipitation revealed that tegaserod treatment reduced the binding between YTHDF1 and *CCNE2* mRNA (Fig. 6h). Moreover, tegaserod inhibited m^6^A-mediated *CCNE2* translation in a luciferase assay (Fig. 6i). Of note, tegaserod treatment did not affect the expression of YTHDF1 (Fig. 6f-g), implying that tegaserod inhibits YTHDF1 function instead of its expression. Collectively, our results indicated that tegaserod directly blocks the binding of YTHDF1 with m^6^A-modified mRNAs, leading to reduced mRNA translation.

### Tegaserod impedes AML progression and prolongs survival

To investigate the potential anti-AML function, we evaluated the effect of tegaserod on AML cells *in vitro*. Tegaserod inhibited THP-1 cell growth in a dose-dependent manner (Fig. 7a). The half-maximal inhibitory concentration (IC50) was 3.58 μM. Similar to YTHDF1 knockdown, tegaserod significantly inhibited cell proliferation, induced G1 phase arrest, and induced cell apoptosis in AML patient-derived CD34^+^ LSCs in a dose-dependent manner (Fig. 7b-d). Moreover, tegaserod inhibited the colony-forming capability of AML patient-derived CD34^+^ LSCs (Fig. 7e and Supplementary Fig. 6a). Tegaserod treatment resulted in a remarkable decrease in the frequency of CD34^+^ LSCs (Fig. 7f).

**Figure 7.**
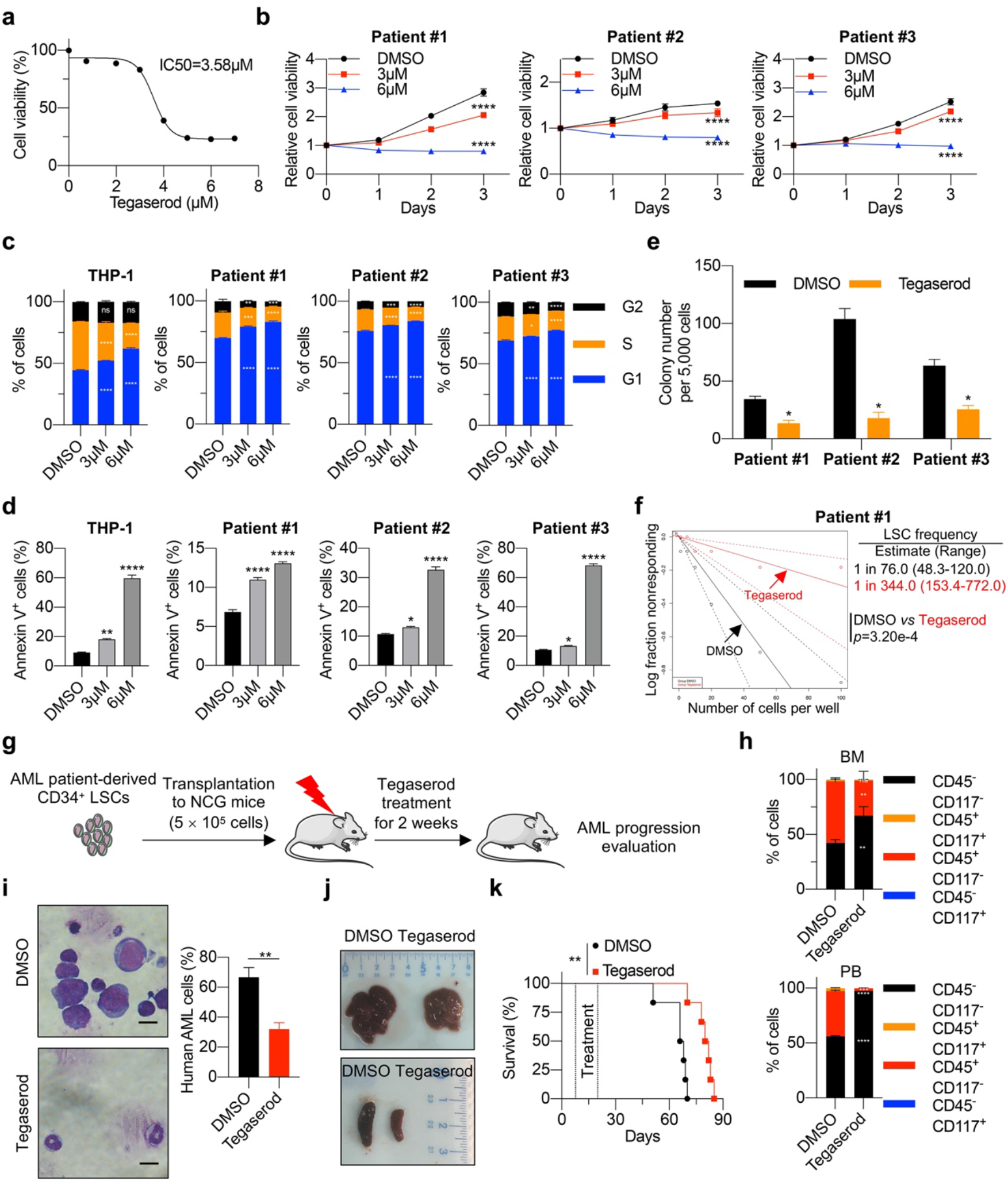
Tegaserod impedes AML progression and prolongs survival. (a) IC50 fitting curve of tegaserod maleate treatment in THP-1 cells for 48 hours. (b) Effect of tegaserod maleate on cell viability of AML patient-derived CD34^+^ LSCs. Data are represented as mean ± SEM (n = 3). **** *p* < 0.0001, two-way ANOVA. (c) Effect of tegaserod maleate on cell cycle of THP-1 cells and AML patient-derived CD34^+^ LSCs. The quantitative data are represented as mean ± SEM (n = 3). * *p* < 0.05, ** *p* < 0.01, \*\*\**p* < 0.001, \*\*\*\**p* < 0.0001, ns, not significant, two-way ANOVA. (d) Effect of tegaserod maleate on apoptosis of THP-1 cells and AML patient-derived CD34^+^ LSCs. The quantitative data are represented as mean ± SEM (n = 3). * *p* < 0.05, ** *p* < 0.01, \*\*\*\**p* < 0.0001, t-test. (e) Colony-forming units of AML patient-derived CD34^+^ LSCs treated with DMSO or tegaserod maleate. The quantitative data are represented as mean ± SEM (n = 3). * *p* < 0.05, t-test. (f) LSC frequency changes in AML patient-derived CD34^+^ LSCs upon tegaserod maleate treatment as estimated by *in vitro* limiting dilution assays (LDAs). (g) Schematic of xenograft mouse model. Irradiated (1 Gy) immunodeficient mice (NCG) were injected with AML patient-derived CD34^+^ LSCs and then treated with tegaserod maleate (2.5 mg/kg) or DMSO every other day for 2 weeks. (h) Percentage of human CD45^+^ and CD117^+^ cells in BM and PB of mice treated with tegaserod maleate or DMSO. The quantitative data are represented as mean ± SEM (n = 3). ** *p* < 0.01, *** *p* < 0.001, **** *p* < 0.0001, ns, not significant, two-way ANOVA. (i) Wright-Giemsa staining of BM from mice treated with tegaserod maleate or DMSO. Data (right panel) are represented as mean ± SEM. ** *p* < 0.01, t-test. Scale bar, 10 μm. (j) Effect of tegaserod maleate on mice hepatosplenomegaly. (k) Survival curve of mice transplanted with AML patient-derived CD34^+^ LSCs with or without tegaserod maleate treatment. Mantel-Cox test, ** *p* < 0.01.

Finally, we performed experiments to assess potential toxicities in mice using 5mg/kg tegaserod every other day for 2 weeks, the results showed that tegaserod did not affect blood, liver, spleen and kidney (Supplementary Fig. 5a-d). Then, we assessed the anti-AML role of tegaserod *in vivo* (Fig. 7g). The mice transplanted with CD34^+^ LSCs were intraperitoneally injected with 2.5 mg/kg of tegaserod every other day for 2 weeks. Tegaserod significantly decreased human AML disease burden and lessen hepatosplenomegaly in xeno-transplanted mice (Fig. 7h-j and Supplementary Fig. 6b-c). Tegaserod reduced leukemia cells infiltration of liver architecture (Supplementary Fig. 6d). Likewise, peripheral blood analysis revealed a reduction in the number of WBCs and recovery of RBCs, HGB, and PLTs in tegaserod-treated mice (Supplementary Fig. 6e). Moreover, tegaserod significantly prolonged the survival of xeno-transplanted mice (Fig. 7k). In summary, our data suggested that tegaserod might be repositioned as a new therapeutic agent for hampering AML progression.

## DISCUSSION

So far, the function of m^6^A reader protein YTHDF1 in leukemogenesis remains unexplored. In this study, we demonstrated the upregulated expression of YTHDF1 protein in human AML and an essential role of YTHDF1 in the progression of AML. YTHDF1 not only sustains cell growth and cell cycle progression but also inhibits cell apoptosis. Depletion of YTHDF1 delays progression and prolongs survival in a human AML cells-derived xenograft AML model. However, knockout of *Ythdf1* gene does not affect normal hematopoiesis in mice, indicating that the function of YTHDF1 relies on its expression as YTHDF1 expression is low in normal myeloid cells but dramatically increased in AML cells. However, it is worth noting that the expression of YTHDF1 is regulated at protein level, but not mRNA level in AML. The available TCGA and GSE public datasets are based on mRNA level, not protein level. Not surprisingly, these datasets and our qPCR data all showed no difference between normal human and AML patients. We therefore concluded that not only the mRNA level but also the protein expression of YTHDF1 should be considered for the clinical relevance of YTHDF1. Taken together, these results demonstrate that YTHDF1 is selectively required for leukemogenesis, providing a potential therapeutic target for AML.

By combining transcriptome-wide m^6^A-seq, RNA-seq, and Ribo-seq, we identified a series of oncogenic pathways positively regulated by YTHDF1. In particular, we have validated CCNE2 as a target of YTHDF1. CCNE2 has been demonstrated as a key regulatory factor in G1/S phase transition in a variety of solid cancers (24). In line with its reported function, genetic ablation of CCNE2 inhibits AML cell proliferation and S phase progression. Of note, overexpression of CCNE2 rescues the phenotype loss in YTHDF1-deficient cells, indicating that CCNE2 contributes to YTHDF1-promoted leukemogenesis. Elevated CCNE2 expression has been reported in AML samples (25). Our data revealed a novel mechanism of *CCNE2* translation involving m^6^A installation and YTHDF1-binding, accounting for the upregulation of CCNE2 in AML. YTHDF1 is well known for its function in promoting the translation initiation of m^6^A-modified mRNAs. It was proposed that YTHDF1 promotes 40S ribosome small subunits loading after recognizing m^6^A located at 3’ UTR (12). However, a majority of YTHDF1-binding m^6^A sites, including the m^6^A sites in *CCNE2* mRNA, lie in the coding region (CDS) of the transcripts, where translation elongation happens. Of note, a recent study demonstrated that YTHDF1 regulates the translation of *SNAIL* mRNA through the m^6^As in CDS (26). Moreover, the interaction between YTHDF1 and elongation factors strongly indicates the involvement of YTHDF1 in translation elongation (26), while the detailed mechanism needs further investigation.

Both YTHDF1 and YTHDF2 play oncogenic roles in AML (27), but specific mechanisms are not the same. Our data showed that YTHDF1 promotes the translation of a subset of oncogenes, whereas YTHDF2 was reported to facilitate the degradation of tumor suppressor mRNAs in AML (27). These data are in line with the current working model that YTHDF1 mainly affects mRNA translation while YTHDF2 regulates mRNA decay (11,12). However, YTHDF1 and YTHDF2 share a large set of common target mRNAs, and they might compete with each other in mRNA-binding (12). Indeed, a recent study reported that YTHDF proteins show identical binding to all m^6^A sites in mRNAs and have a certain overlap in their roles under certain circumstances (28). These seemingly contractionary results strongly suggest context-dependent functions of YTHDF family members (29). Additional factors, such as binding proteins of YTHDF members, and other RNA-binding proteins, should participate in the regulation of the function of YTHDF proteins and work cooperatively to determine the final fate of m^6^A-modified mRNAs.

To date, the drugs targeting YTHDF1 have not been discovered. In this study, we tried to screen for candidate inhibitors of YTHDF1 from FDA-approved drugs. Fortunately, tegaserod, an approved drug for the treatment of irritable bowel syndrome with constipation (18), was predicted to target the YTH domain of YTHDF1. Tegaserod maleate was voluntarily withdrawn in 2007 due to possible cardiovascular events, and reintroduced in the United States in 2018 (30). We found that tegaserod showed inhibitory effect on the binding of the YTH domain with m^6^A-modified mRNAs in a concentration-dependent manner, and the inhibited rate was more than 20%. However, tegaserod has little effect on the binding of m^6^A/YTHDF2 or m^6^A/YTHDF3. We proposed that although some key residues are conserved, the YTH domains of YTHDF1/2/3 are still different. Since we used the YTH domain of YTHDF1 for the drug screening, tegaserod specifically recognizes YTHDF1 instead of DF2 or DF3. Moreover, tegaserod inhibited the translation of *CCNE2* mRNA in a similar pattern as YTHDF1 knockdown. Based on these results, tegaserod might serve as an inhibitor for future biological studies on YTHDF1. Importantly, tegaserod has drawn great attention recently for its potential role in cancer treatment, including lung cancer and melanoma(31,32). Our data showed that tegaserod could inhibit AML cell proliferation *in vitro* and AML progression in a human AML cells-derived xenograft model, indicating tegaserod as a candidate therapeutic inhibitor for AML.

In conclusion, we highlighted the functional importance of YTHDF1 in promoting AML progression through regulating mRNA translation in an m^6^A-dependent manner. We also provided strong evidence for repositioning tegaserod as an anti-leukemia drug targeting YTHDF1. Given that YTHDF1 is dysregulated in a wide spectrum of cancers (21,33,34), additional studies in other cancer types will identify whether targeting YTHDF1-regulated gene expression can be broadly applied for cancer treatment.

## METHODS

### Mouse experiments

Generation of *Ythdf1* conditional knockout (*Ythdf1^cKO^*) mice was described(21), *Ythdf1^+/-^* mice were generated by mating *Ythdf1^fl/fl^* mice with CMV-Cre mice. *Ythdf1^-/-^* were generated by inter-cross *Ythdf1^+/-^* mice. Mice were maintained and bred in specific pathogen-free conditions at the Animal Center of Zhejiang University. All animal studies were performed in compliance with the Guide for the Care and Use of Laboratory Animals by the Medical Experimental Animal Care Commission of Zhejiang University. All animal studies used the protocol that has been approved by the Medical Experimental Animal Care Commission of Zhejiang University.

### Cell culture

THP-1 and MV-4-11 human leukemia cells were obtained from ATCC. THP-1 cells were cultured in RPMI Medium 1640 basic (Gibco) supplemented with 10% fetal bovine serum (Thermo Fisher Scientific) and 1mM sodium pyruvate (Gibco). MV-4-11 cells were cultured in Iscove’s Modified Dulbecco’s Medium (Gibco) supplemented with 10% fetal bovine serum (Thermo Fisher Scientific). AML patient-derived CD34^+^ leukemia stem cells (LSCs) were cultured in StemSpan SFEM (StemCell Technologies) supplemented with rhIL-3 (50 ng/mL), rhFLT3-Ligand (50 ng/mL), rhTPO (50 ng/mL), and rhSCF (50 ng/mL) (Peprotech). Cells were maintained at 37 °C in an atmosphere containing 5% CO_2_.

### Leukemia patients and healthy donors’ samples

The leukemic samples were obtained at the time of diagnosis and with informed consent at Central People’s Hospital of Zhanjiang. The non-leukemic controls samples from healthy donors and normal cord blood units were collected and with informed consent at Central People’s Hospital of Zhanjiang. The study was approved by the Medical Ethics Committee of Central People’s Hospital of Zhanjiang and conducted in accordance with the Declaration of Helsinki. AML CD34^+^ leukemia stem cells (LSCs) were isolated using CD34^+^ beads (Miltenyi Biotec).

### Flow cytometry analysis

For intracellular staining of YTHDF1 in leukemic samples, PB were lyzed in RBC lysis buffer, fixed and permeabilized, incubated with primary antibody-YTHDF1, followed by incubation with the secondary antibody.

For mice, BM cells were isolated by washing tibias and femurs cut off both ends of the epiphysis using PBS with a 1 mL syringe. For the expression of hematopoietic stem cells markers analysis, unfractionated BM cells were stained with anti-c-Kit (CD117) and anti-Sca-1 antibodies. For analysis of differentiated cells, PB were stained with anti-CD3, anti-CD4 and anti-CD19 antibodies for T and B cells; Anti-CD11b and anti-Gr-1 antibodies for myeloid cells, lyzed in RBC lysis buffer.

Leukemia cells with different treatments were harvested and washed with PBS, to analyze cells undergoing apoptosis, cells were suspended in binding buffer containing Annexin V and Propidium iodide.

Flow cytometry analysis were performed using FACS Canto II flow cytometer (BD Biosciences) and ACEA NovoCyteTM (ACEA Biosciences). The antibodies used for flow cytometry were listed in Supplementary Table 5.

### Cell cycle analysis

For cell cycle analysis, cells were collected after treatment, fixed in 70% ethanol for overnight, and stained with PI (40 μg/ml, Beyotime) plus 0.2 mg/ml of RNase A (Thermo Fisher Scientific) for 1 hour at 4 °C.

### Histology and slides

Livers and spleens were rinsed with PBS, fixed in 10% formalin, paraffin-embedded and sectioned at 5 mm. Sections were stained with hematoxylin and eosin (H&E). Smear slides from PB and BM were let dry and stained with Wright-Giemsa Stain Solution and mounted with Poly-mount.

### *In vitro* colony-forming assay and limiting dilution assays

Cells were plated in methylcellulose medium (StemCell Technologies) according to the manufacturer’s instructions. Colonies were evaluated after 14 days of incubation. For *in vitro* limiting dilution assays (LDAs), cells were suspended in methylcellulose medium and plated in 96-well plates at a limiting dilution manner, for each dose, 12 wells were included. The number of wells containing spherical colonies was counted after 10 days. ELDA software was used to estimate the frequency of LSC (35).

### Xenotransplantation experiments

To determine the effect of YTHDF1 knockdown on the leukemia-initiating capacity of AML patient-derived CD34^+^ LSCs, 8-week-old male NOD CRISPR *Prkdc Il2r Gamma* (NCG) mice were sub-lethally irradiated (2 Gy of X-ray) and injected with the indicated cells via tail vein (1 × 10^6^ cells per mouse).

For tegaserod treatment, 8-week-old NCG mice were irradiated with 1 Gy of X-ray and injected with AML patient-derived CD34^+^ LSCs (5 × 10^5^ cells per mouse). On the seventh-day post-transplantation, tegaserod maleate (2.5 mg/kg) in DMSO or DMSO only was intraperitoneally injected into mice every other day for 2 weeks.

### Virtual screening and molecular docking

The structure of the YTH domain of YTHDF1 was downloaded from the PDB database (http://www.rcsb.org), and prepared using protein preparation wizard (Glide, Schrödinger, LLC, New York, NY, USA). After removing the crystallographic waters and the unnecessary chain, the missing hydrogens were added. Then, the overall protein was sent to minimize using OPLS 2005 force filed with a default constraint of 0.30 Å RMSD (Root Mean Square Deviation). The grid box was generated by centered on residue D401 (20 Å × 20Å × 20Å) in the receptor-grid generation module. The molecular docking of 1805 FDA-approved drugs was performed using GLIDE. Firstly, all the prepared compounds were docked using the HTVS (High Throughput Virtual Screening) mode to estimate protein-ligand binding affinities. Then, standard precision (SP) docking was performed to redock the top ranked compounds from HTVS. Finally, the top ranked compounds were subjected to another round of docking by using extra precision (XP) mode.

### Cloning, Expression, and Purification of the C-termus of YTHDF1, YTHDF2 and YTHDF3

The coding sequence of YTHDF1 C-terminus containing YTH domain (amino acids 361-559), YTHDF2 C-terminus containing YTH domain (amino acids 401–554) and YTHDF3 C-terminus containing YTH domain (amino acids 407–560) was cloned into a modified pET28b vector, which contains an N-terminus 6× Histidine SUMO tag. Recombinant fusion protein was overexpressed at 18 °C in the E. coli strain Rosetta (Novagen) and purified by Ni-NTA Agarose Resin (Yeasen Biotech). The His-sumo tag was cleaved by ULP1 and subsequently removed by a second step Ni-NTA Agarose Resin purification. YTH domain was further purified with a cation-exchange column (HiTrap SP FF) and a Superdex 75 increase column on AKTA Pure (GE Healthcare).

### Surface plasmon resonance (SPR) assay

Biacore T200 instruments (GE Healthcare) were used to evaluate the binding affinity of m^6^A RNA or non-m^6^A RNA with the YTH domain of YTHDF1. Briefly, 20 nmol/L of biotinylated m^6^A RNA oligos or non-m^6^A RNA oligos was captured on the surface of the StrepAvidin chip at a flow rate of 30 μL/min in phosphate buffered saline (PBS) with 0.05% (v/v) Tween-20 and 5% DMSO, pH 7.4. The oligo sequences were listed in Supplementary Table 6. The indicated concentration of the YTH domain with or without tegaserod was injected into the flow system and analyzed for 90 s, and the dissociation was 120 s. All binding analysis was performed in phosphate buffered saline with 0.05% (v/v) Tween-20 and 5% DMSO, pH 7.4, at 25 °C. The association time was set to 120 s, while the dissociation time was set to 360s. After dissociation, the chip surface was regenerated by 50 mM NaOH and 1M NaCl. Prior to analysis, double reference subtractions were made to eliminate bulk refractive index changes, injection noise, and data drift. The binding affinity was determined by global fitting to a Langmuir 1:1 binding model within the Biacore Evaluation software (GE Healthcare).

### Statistical analysis

Statistical analysis was performed using GraphPad Prism 8 software (GraphPad Software, Inc.). Asterisks denote statistical significance (ns, not significant; * *p* < 0.05; \*\**p* < 0.01; *** *p* < 0.001; **** *p* < 0.0001).

### Other methods

Cell viability analysis, plasmids construction, immunoblotting, qPCR, luciferase assay, M^6^A RIP, YTHDF1 RIP, RNA-seq, and Ribosome profiling were performed as described previously (21). The detailed methods were described in the supplementary data. The primary antibodies used were listed in Supplementary Table 5. The primers used for the cloning were listed in Supplementary Table 6.

## ACKNOWLEDGEMENTS

We thank doctors Yuming Zhang, Honghua He, Liang Liang, Lirong Nie, Qinghua Li, Wenjun Liang, Jie Long, and Jingzhi Ma at the Affiliated Hospital of Guangdong Medical University for assisting in collecting the clinical samples. We thank doctors Yi-Hong Xiao and Yanqing Lin at the Central People’s Hospital of Zhanjiang for assisting in collecting the clinical samples. This work was supported by grants from the National Natural Science Foundation of China (81971886 to Z.C. and 81672847, 82073110 to X.G.), the Zhujiang Talent Program (2019QN01Y279), and the Basic and Applied Basic Research Fund of Guangdong Province (2020A1515010468).

## AUTHOR CONTRIBUTIONS

Z.C. and X.G. conceived the project and designed the experiments. Y.H. performed the majority of the experiments. Y.C. and T.L. performed flow cytometry analysis. Y.Z. assisted the molecular experiments. Z.Y., G.W., Y.C., J.X., R.W., W.L., Y.Z. and J.C. provided the clinical samples. Y.H. and X.G. wrote the manuscript. All authors discussed the results and commented on the manuscript.

## DATA AVAILABILITY

The data that support the findings of this study are available from the corresponding author upon reasonable request. The sequencing data including m6A-seq, Ribo-seq, and RNA-Seq data, are available in Gene Expression Omnibus under accession number GSE184842 (https://www.ncbi.nlm.nih.gov/geo/query/acc.cgi?acc=GSE184842).

