## supplemental Figure1-6 for "The RNA m^6^A reader YTHDF1 promotes hematopoietic malignancy by maintaining oncogenic translation"

[#] These authors contributed equally.

### SUPPLEMENTARY INFORMATION

### Supplementary Figures 1-6

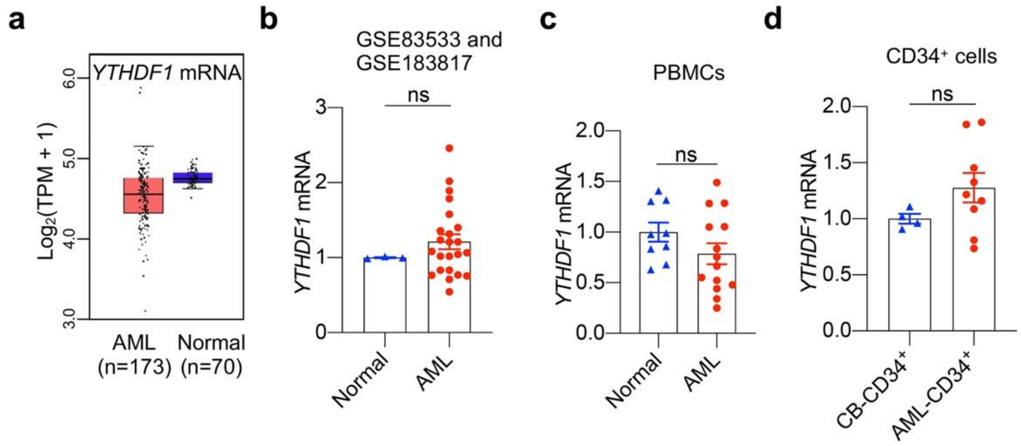


**Supplementary Figure 1. The expression of *YTHDF1* mRNA is not upregulated in AML.**

(a) Box plot showing the relative expression of *YTHDF1* mRNA in normal and AML samples from the TCGA database.

(b) Comparison of *YTHDF1* mRNA expression in healthy donors (n=3) vs primary AML patients (n=23) based on GEO datasets (GSE83533 and GSE183817). The quantitative data are represented as mean ± SEM. ns, not significant, t-test.

(c) Relative *YTHDF1* mRNA expression in PBMCs from Normal (n=9) and human AML patients (n=14). Data are represented as mean ± SEM. ns, not significant, t-test.

(d) Relative *YTHDF1* mRNA expression in normal cord blood-derived $CD34^+$ cells (n = 4) and AML $CD34^+$ LSCs (n = 9). Data are represented as mean ± SEM. ns, not significant, t-test.

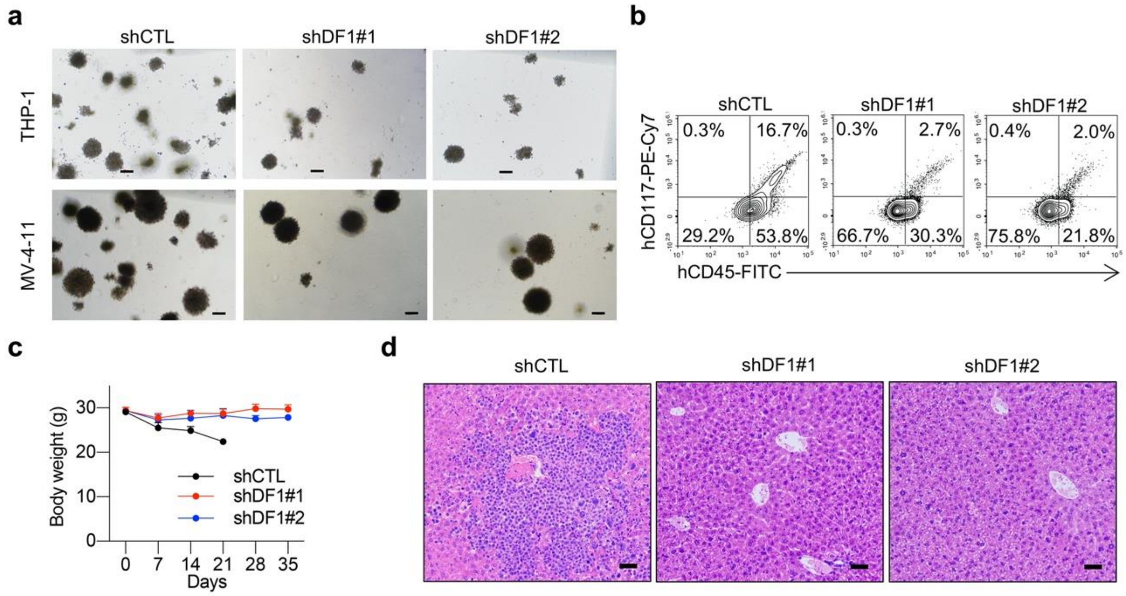


**Supplementary Figure 2. YTHDF1 is essential for the progression of AML.**

(a) Representative pictures of colonies from control and YTHDF1 knockdown THP-1 and MV-4-11 cells as described in Figure 3e-f. Scale bar, 200 μm

(b-d) NCG mice were injected with control or YTHDF1 knockdown AML patient-derived $CD34^+$ LSCs and analyzed as described in Figure 3m.

(b) Flow cytometry analysis of the percentage of human AML cells (human $CD45^+$ and $CD117^+$ cells) in BM of recipient mice as described in Figure 3n.

(c) Body weight of recipient mice was demonstrated.

(d) Hematoxylin and eosin (H&E) staining of liver from recipient mice. Scale bar, 50 μm.

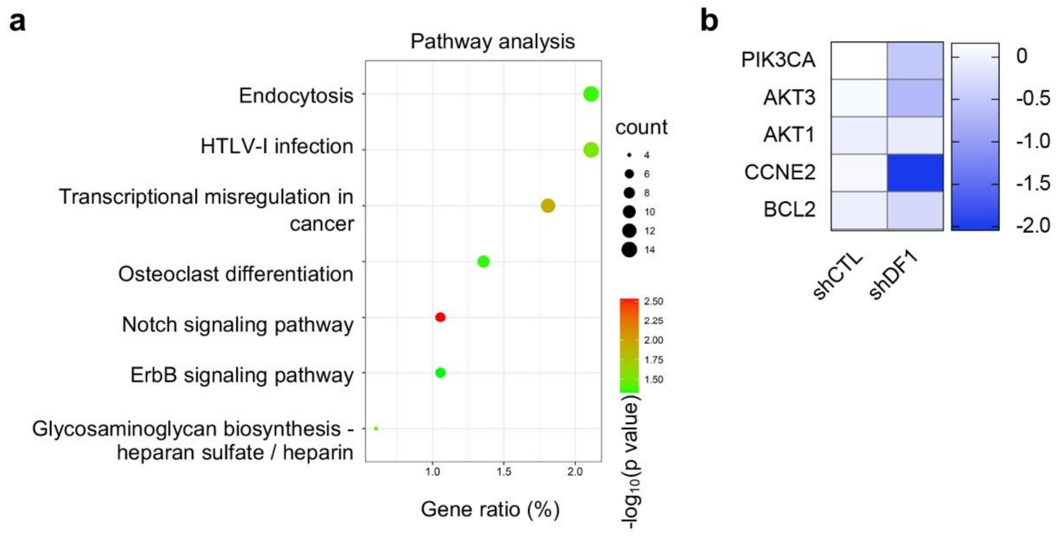


**Supplementary Figure 3. Transcriptome-wide identification of YTHDF1-regulated transcripts.**

(a) KEGG enrichment analysis of YTHDF1 targets that are upregulated in YTHDF1-deficient cells.

(b) Heatmap showing the translational efficiency (TE) of PI3K-Akt signaling components in control and YTHDF1 knockdown cells. The values show the fold change of the indicated color of the heatmap.

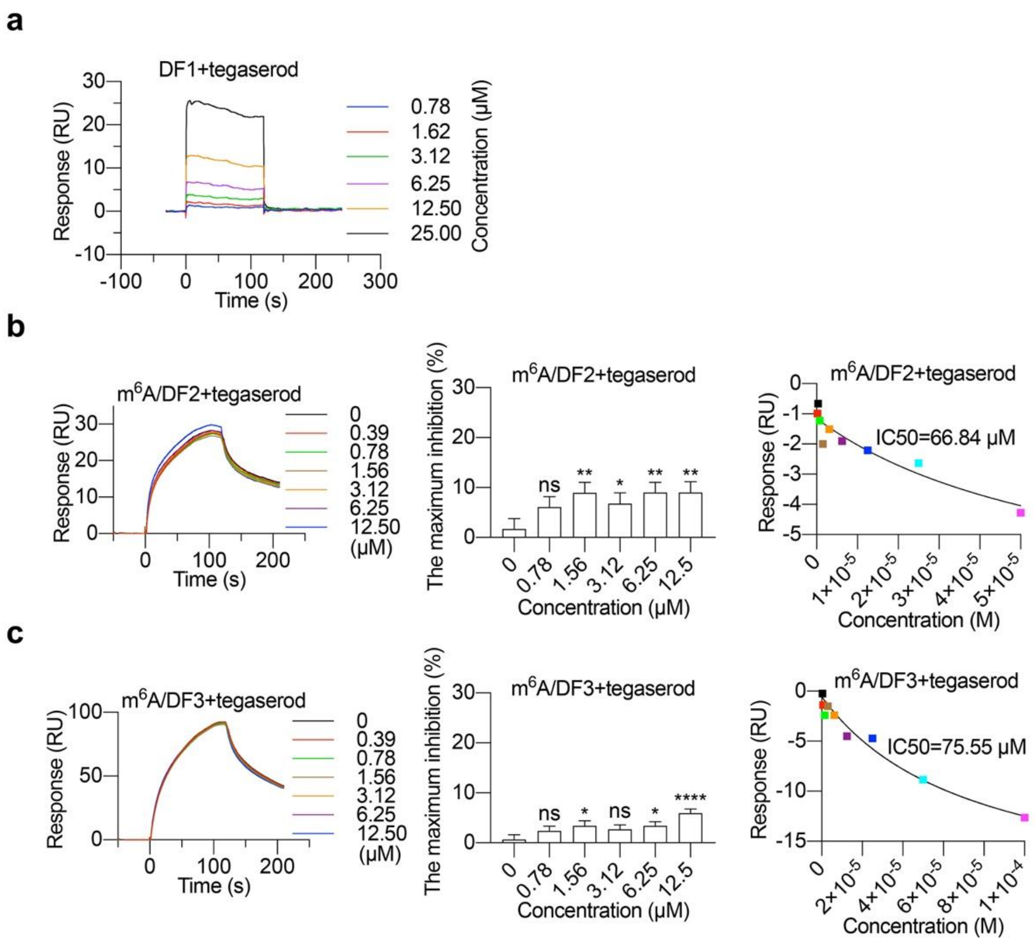


**Supplementary Figure 4. The specificity of tegaserod on m$^6$A/YTHDF1 binding.**

(a) Biacore assay of the binding between YTH domain of YTHDF1 and tegaserod.

(b) Effect of tegaserod on the interaction between m$^6$A-modified oligos and YTH domain of YTHDF2. * $p < 0.05$, ** $p < 0.01$, ns, not significant, t-test.

(c) Effect of tegaserod on the interaction between m$^6$A-modified oligos and YTH domain of YTHDF3. * $p < 0.05$, **** $p < 0.0001$, ns, not significant, t-test.

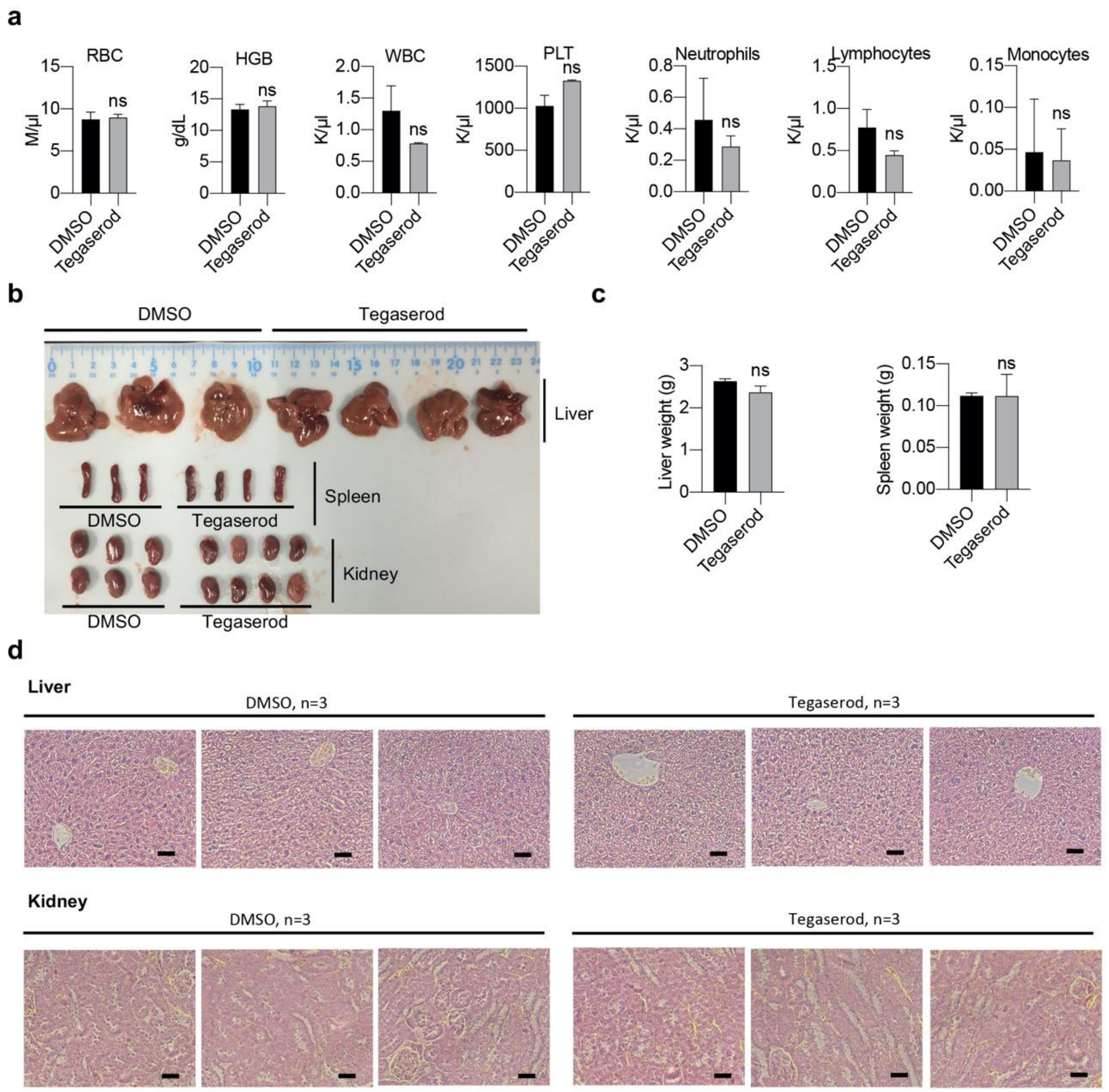


**Supplementary Figure 5. The potential toxicities of evaluating tegaserod in *vivo*.**

(a) Peripheral blood counts from mice treated with DMSO or tegaserod maleate. Data are represented as mean ± SEM (n = 3), ns, not significant, t-test.

(b-d) Effect of tegaserod maleate on liver, spleen and kidney from mice. Data are represented as mean ± SEM (n = 3), ns, not significant, t-test. Scale bar, 50 μm.

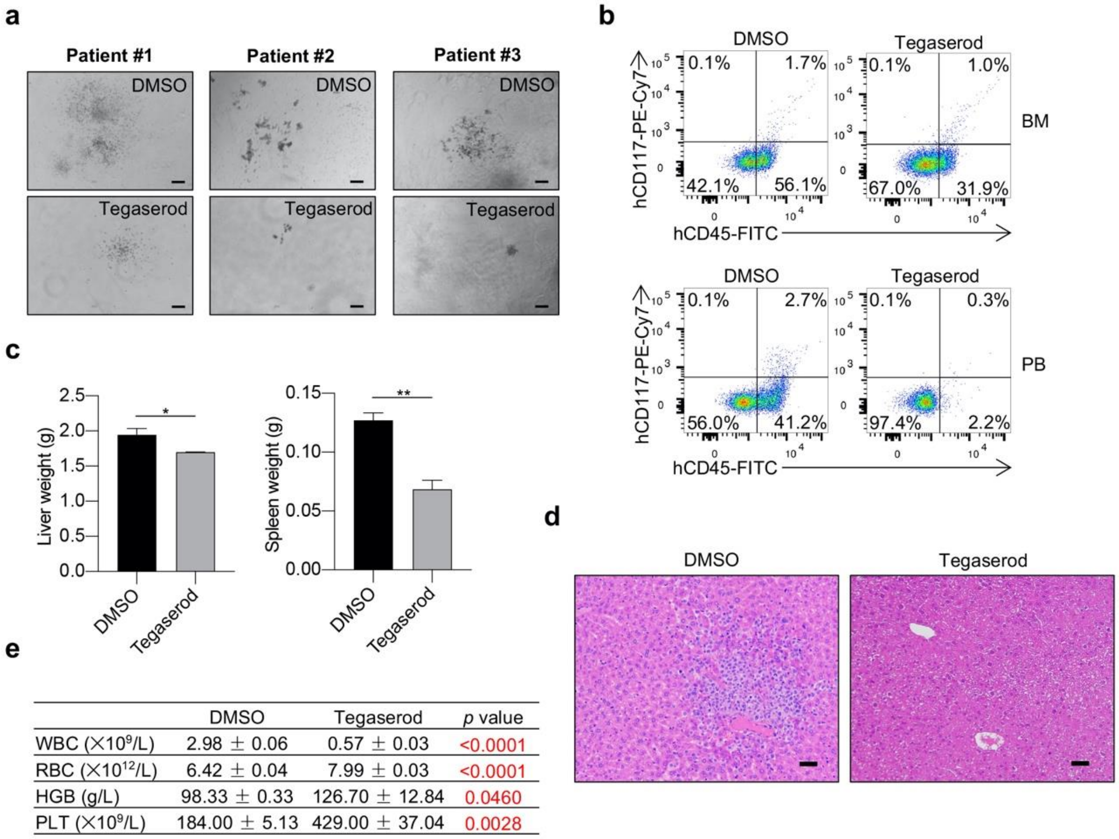


| | DMSO | Tegaserod | *p* value |
|---|---|---|---|
| WBC (×10⁹/L) | 2.98 ± 0.06 | 0.57 ± 0.03 | <0.0001 |
| RBC (×10¹²/L) | 6.42 ± 0.04 | 7.99 ± 0.03 | <0.0001 |
| HGB (g/L) | 98.33 ± 0.33 | 126.70 ± 12.84 | 0.0460 |
| PLT (×10⁹/L) | 184.00 ± 5.13 | 429.00 ± 37.04 | 0.0028 |

**Supplementary Figure 6. Tegaserod impedes AML progression and prolongs survival.**

(a) Representative pictures of colonies from AML patient-derived $CD34^+$ LSCs treated with DMSO or tegaserod maleate as described in Figure 7e. Scale bar, 200 μm

(b) Flow cytometry analysis of the percentage of human AML cells (human $CD45^+$ and $CD117^+$ cells) in BM and PB of mice treated with tegaserod maleate or DMSO as described in Figure 7h.

(c) The weight of liver and spleen from AML patient-derived $CD34^+$ LSCs-transplanted mice treated with DMSO or tegaserod maleate. Data are represented as mean ± SEM (n = 3). * $p < 0.05$, ** $p < 0.01$, t-test.

(d) H&E staining of liver from AML patient-derived $CD34^+$ LSCs-transplanted mice treated with DMSO or tegaserod maleate. Scale bar, 50 μm.

(e) PB counts from AML patient-derived $CD34^+$ LSCs-transplanted mice treated with DMSO or tegaserod maleate. Data are represented as mean ± SEM (n = 3, t-test).
